# Real-Time Spatiotemporal Measurement of Extracellular Signaling Molecules Using an Aptamer Switch-Conjugated Hydrogel Matrix

**DOI:** 10.1101/2023.07.09.548040

**Authors:** Chan Ho Park, Ian A. P. Thompson, Sharon S. Newman, Linus A. Hein, Xizhen Lian, Kaiyu Fu, Jing Pan, Michael Eisenstein, H. Tom Soh

## Abstract

Cells rely on secreted signaling molecules to coordinate essential biological functions including development, metabolism, and immunity. Unfortunately, such signaling processes remain difficult to measure with sufficient chemical specificity and temporal resolution. To address this need, we have developed an aptamer-conjugated hydrogel matrix that enables continuous fluorescent measurement of specific secreted analytes – in two dimensions, in real-time. As a proof of concept, we performed real-time imaging of *Dictyostelium discoideum* cells, a well-studied amoeba model wherein inter-cellular communication is performed though cAMP signaling. We engineered a set of aptamer switches that generate a rapid and reversible change in fluorescence in response to cAMP signals. By combining multiple switches with different dynamic ranges, we can measure cAMP concentrations spanning three orders of magnitude in a single experiment. These sensors are embedded within a biocompatible hydrogel on which cells are cultured and their cAMP secretions can be imaged using fluorescent microscopy. Using this aptamer-hydrogel material system, we achieved the first direct measurements of oscillatory cAMP signaling that correlate closely with previous indirect measurements. Using different aptamer switches, this approach could be generalized for measuring other secreted molecules to directly visualize diverse extracellular signaling processes and the biological effects that they trigger in recipient cells.

## Introduction

Cellular imaging tools such as fluorescent proteins^[1]^, incorporation of non-natural fluorescent amino acids^[2,3]^, and biorthogonal chemistry^[4,5]^ have revolutionized the study of biology, enabling researchers to perform high-resolution visualization of structures and processes within cells or at the surface of the cell membrane. However, many critical biological functions are also coordinated outside of the cell—most notably, cell-to-cell communication, which is typically achieved through the release and diffusion of secreted signaling molecules. Molecular communication between cells governs many important phenomena, ranging from the collective behavior of relatively simple organisms in bacterial biofilms^[6]^ to complex processes in human physiology like neurotransmission^[7]^. Current techniques that correlate cellular behavior with the dynamics of molecular signals inside cells provide clear evidence for the importance and spatiotemporal complexity of extracellular communication^[8]^. However, these methods cannot directly detect and monitor the extracellular signals themselves.

Cell culture matrices that would allow us to record such extracellular signaling processes directly and quantitatively as real-time ‘molecular movies’ would be invaluable for directly identifying and investigating the role of signaling molecules in coordinating diverse cellular behaviors. This requires a sensing material with high sensitivity and molecular specificity to distinguish molecules of interest from the wide range of interferents present in the extracellular milieu. Furthermore, detection must occur rapidly and reversibly to ensure accurate measurement of signals that may be rapidly released and then degraded or re-absorbed. Finally, measurements must take the form of an imaging method that is able to capture a broad field of view with high spatial resolution to capture both local complexity and phenomena that are coordinated across considerable distances within tissues or other multicellular assemblies. Various groups have made progress in tackling these technical challenges, beginning with optical interferometry-based methods that can detect protein secretion^[9]^. Unfortunately, this technique lacks the chemical specificity to discriminate between proteins of similar size. Subsequent approaches have achieved improved specificity by binding secreted proteins of interest with affinity reagents and mapping their distribution with refractive-index-based optical measurements^[10–13]^. However, these methods are vulnerable to nonspecific binding by the affinity reagents to the various interferents present in complex media, and are generally applicable only to relatively large analytes—typically proteins. This is an impediment to the analysis of cellular communications mediated by small molecules. Other groups have used binding-induced near-infrared fluorescence shift in aptamer-functionalized single-walled carbon nanotubes to track protein and small molecule secretions^[14,15]^. However, the images generated by these methods are sparsely labeled and thus cannot capture the rich detail of extracellular signaling across the full field of view.

In this work, we present a materials platform that uses fluorescent aptamer switches to achieve rapid, label-free, and spatially resolved imaging of extracellular communication via specific small-molecule signals in real time. Our SEMAPHORE (SEcretion Mapping through APtamer-Hydrogels with Optical REporting) platform embeds aptamer switches that produce a fluorescent response to target binding within a thin layer of biocompatible polyethylene glycol (PEG)-based hydrogel matrix. Cells can be cultured on this hydrogel surface, and as cell-to-cell communication occurs, secreted biomolecules diffuse into the hydrogel, reversibly activating a localized fluorescent response. The hydrogel can be continuously imaged using standard fluorescent microscopy, enabling real-time measurement of the release and diffusion of these molecular signals across a wide 2D field of view. We can achieve accurate and quantitative molecular measurements by tuning the aptamer switch response for target sensitivity across a broad concentration range, with rapid switching kinetics that capture the dynamics of analyte release and reuptake. As proof of concept, we used SEMAPHORE to continuously monitor the dynamics of cyclic adenosine 3’,5’-monophosphate (cAMP) secretion during collective migration of *Dictyostelium discoideum* (*Dictyostelium*) cells. We provide a direct measurement of the extracellular cAMP oscillatory behavior that drives *Dictyostelium* cell migration during starvation, obtaining results that match with prior observations based on intracellular reporters and single-timepoint secretion measurements. This new tool for studying cellular communication should be broadly adaptable to new cellular systems by generating aptamer switches that selectively respond to other secreted analytes.

## Results

### The SEMAPHORE Platform

SEMAPHORE comprises a biocompatible hydrogel matrix functionalized with aptamer switches that serve as fluorescent optical sensors to spatiotemporally resolve secreted biomolecules with high specificity and fast and reversible signaling (**Figure 1A**). Cultured cells can be transferred to the surface of SEMAPHORE with minimal disruption of their biological function, and then exposed to a range of stimuli. Signaling biomolecules secreted by the cells diffuse into the hydrogel, where they interact with the embedded aptamer switches. These aptamer switches undergo reversible conformational switching in response to target binding, generating a fluorescent signal that is localized to the immediate proximity of the secreted molecule. Thus, one can continuously measure the hydrogel response using fluorescent microscopy to obtain real-time, spatial quantification of secreted signaling molecules.

**Figure 1.**
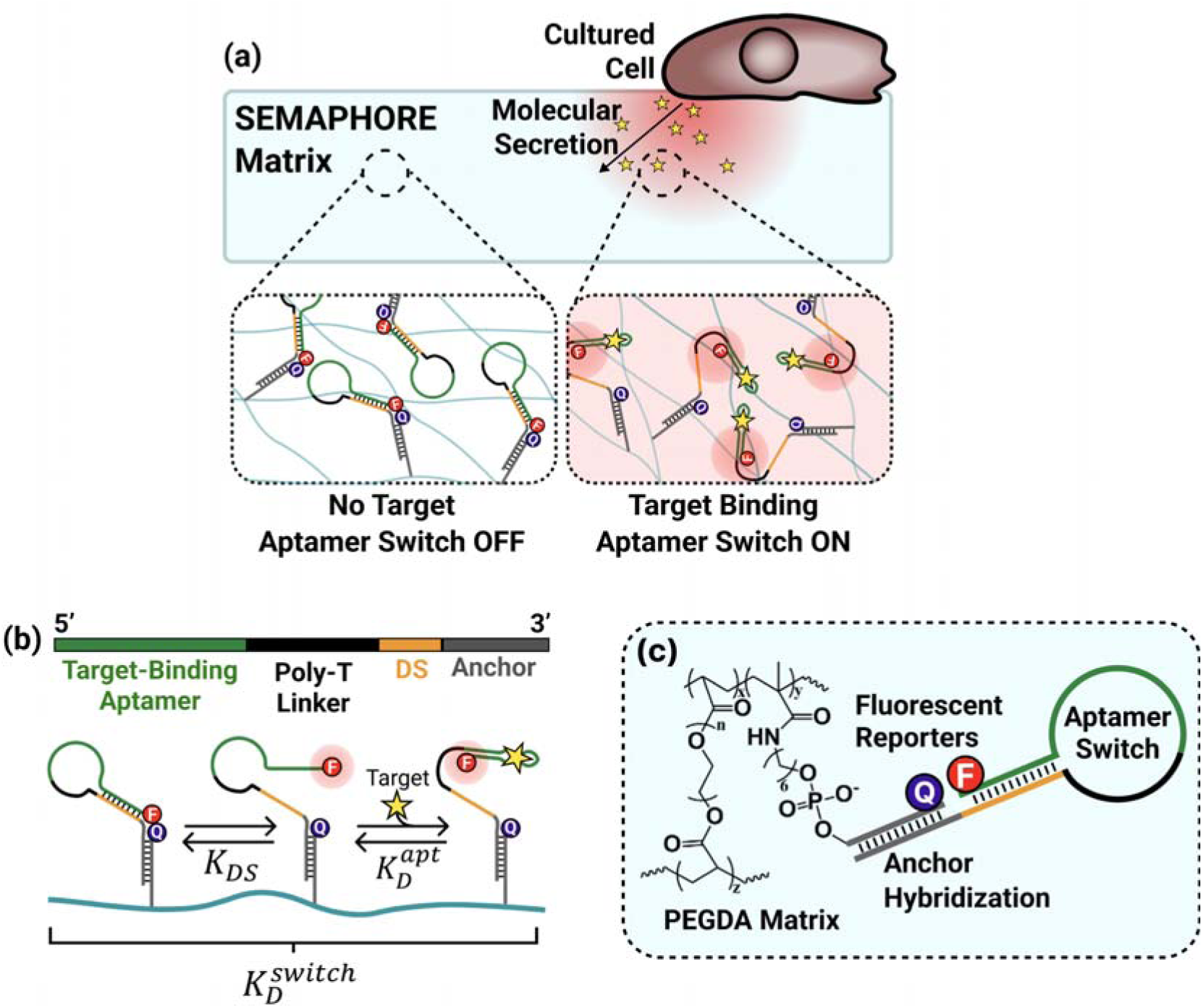
Real-time mapping of signaling molecules using SEMAPHORE. (A) Cells are cultured directly on the SEMAPHORE matrix. Secreted signaling molecules diffuse into the hydrogel, where they bind target-specific aptamer switches, leading to localized fluorescent signaling. (B) Schematic of the intramolecular strand displacement (ISD) aptamer switches used for cAMP detection. The aptamer is connected to a displacement strand (DS) domain via a linker sequence, with a 3’ anchor sequence for immobilization onto a complementary anchor strand incorporated into the hydrogel. Fluorescent switching is achieved when target binding competitively displaces the DS from the aptamer, thereby changing the proximity of a fluorophore-quencher pair within the switch. (C) The SEMAPHORE platform consists of a poly(ethylene glycol) diacrylate (PEGDA) matrix functionalized with fluorescent aptamer switches through hybridization to anchor DNA strands covalently linked to the matrix.

We chose DNA aptamer switches as molecular sensors because they are capable of rapid binding-induced signaling and can be readily coupled to the hydrogel matrix in a chemically defined manner^[16]^. Aptamer switches can be engineered in a variety of ways to achieve fluorescent sensing of a wide range of biomolecules^[17–19]^. We opted to use an intramolecular strand displacement (ISD) design strategy in which the aptamer is linked to a displacement strand (DS) sequence through a tunable poly-T linker (**Figure 1C**). The DS hybridizes to the aptamer binding pocket in the absence of target, bringing a fluorophore and quencher into close proximity with each other. Aptamer-target binding competitively disrupts DS hybridization, causing the ISD structure to switch to an open state, which increases the distance between the fluorophore and quencher moieties, leading to enhanced fluorescent signal. The overall affinity of this switch 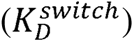 depends on competition between aptamer-target binding (defined by the constant 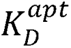) and intramolecular DS hybridization (defined by the constant *K_DS_*). By tuning the lengths of the linker and DS domain, one can therefore optimize the sensitivity of the switch to different target concentrations. Importantly, this switching response is reversible, such that the localized signals will dissipate as secreted concentrations drop and the analyte diffuses away from a given region of the hydrogel, making it possible to continuously track changing target concentrations. These aptamer switches are immobilized within the hydrogel matrix via an anchor DNA sequence appended to the 3’ end of the aptamer switch, which is designed to hybridize to a complementary anchor sequence covalently linked within the hydrogel scaffold (**Figure 1B**). Fluorescent readout of aptamer switching is achieved by labeling the 3’ end of the aptamer and 5’ end of the hydrogel anchor with a fluorophore (Cy3) and quencher (BHQ2), respectively.

We used poly(ethylene glycol) diacrylate (PEGDA) as the main matrix for the hydrogel due to its hydrophilicity, neutral charge in aqueous solutions, and lack of chemical reactivity with cells. The hydrogel scaffold was synthesized and coupled to the BHQ2-labeled anchor DNA strands through 5LJ-end acrydite modifications using a controlled photopolymerization reaction (**Figure S1**). We chose to polymerize the quencher-bearing strand instead of the fluorophore-bearing strand because of the quencher’s greater stability under the UV light conditions used for hydrogel polymerization. By limiting the duration of polymerization, we could minimize UV-induced damage to the DNA strands and BHQ2 quencher (**Figure S2**). Subsequently, dye-labeled aptamers were coupled to the newly synthesized hydrogel via hybridization to the anchor strand.

### Engineering SEMAPHORE to sense cAMP secretions in migrating *Dictyostelium* cells

As a proof of principle, we designed a SEMAPHORE system to record extracellular cAMP and used the social amoebae *Dictyostelium,* a widely used and well-characterized model system for the study of chemotaxis and cell fate decisions^[20,21]^. As *Dictyostelium* cells undergo starvation, they secrete, sense, and respond to cAMP, giving rise to waves of cAMP release that are essential to drive a distinctive morphogenetic cycle of development that starts with the directed migration of the cells towards cAMP signals and the formation of aggregates that calumniate into the formation of fruiting bodies composed of spores atop a stalk of vacuolated cells^[22,23]^.

We constructed our switches from an adenosine-binding aptamer that is well-characterized both in terms of its cAMP-binding properties and its structure^[24]^. As noted above, the ISD design offers the capacity to modulate its target-binding properties based on changes to the DS and linker lengths, and we assessed this tunability by designing cAMP-responsive ISD switches with a wide range of affinities. Specifically, we connected the aptamer to a 7-nt long DS domain through poly-T linkers with lengths varying from 5–33 nt (see **Table S1** for all sequences used in this work). We then measured the affinity of these various switches by immobilizing them within the SEMAPHORE hydrogel and exposing the hydrogel to different cAMP concentrations in buffer (**Figure 2A**). As the length of the linker increases, the confinement between the DS and aptamer is weakened, thereby weakening its competition with aptamer binding (higher *K_DS_*) and increasing the aptamer’s target-binding affinity (lower 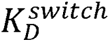). Accordingly, an ISD with a very short 5-nt linker (construct T5) showed low cAMP affinity 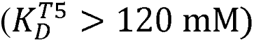, with a limit of detection (LOD) of 247 μM, making it best suited for achieving cAMP detection at very high concentrations. Even slightly lengthening the linker from 5 nt to 9 nt (T9) resulted in a 540-fold jump in cAMP affinity relative to the T5 construct (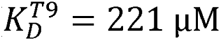, LOD = 80 µM). When we increased the linker length all the way from 5 to 33 nt (construct T33), this increased the affinity by over 1,800-fold, yielding a 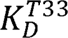 of 66 µM and an LOD of 2.2 μM.

**Figure 2.**
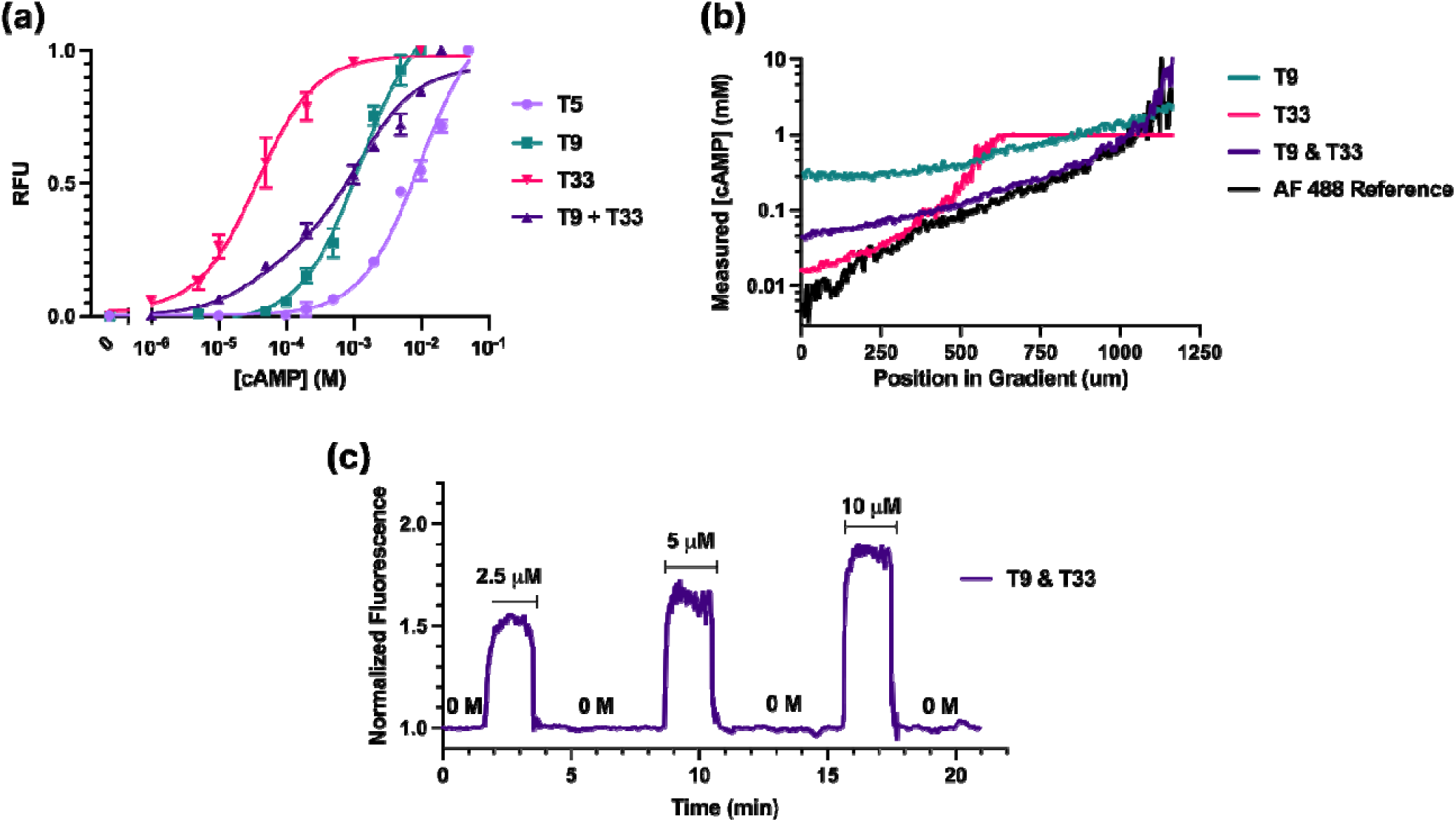
Tuning the binding properties of aptamer switches to optimize SEMAPHORE performance. (A) Fluorescence response of the SEMAPHORE system containing cAMP-responsive aptamer switches with varying poly-T linker lengths (*e.g.*, T5 denotes a 5-nucleotide poly-T linker). Combining the T9 and T33 switches at a 1:1 ratio within the same hydrogel enables sensing over a broader range of cAMP concentrations. (B) Spatial measurement of an exogenously applied cAMP gradient. After measuring the fluorescent signal, cAMP concentrations were estimated by using the responses from panel A as calibration curves. To assess the accuracy of SEMAPHORE, we also directly measured a fluorescent dye (AF 488 Reference) with similar diffusion properties to cAMP under the same gradient conditions. (C) Time-resolved measurement of the SEMAPHORE fluorescent response when exposed to varying pulses of cAMP shows rapid and reversible detection of cAMP across variou concentrations.

Sensors using a single receptor can generally provide accurate molecular quantification over a ∼100-fold concentration dynamic range^[25]^, but an even broader range is necessary to capture the possible range of cAMP dynamics within *Dictyostelium* populations. Average extracellular cAMP concentrations within chemotactic *Dictyostelium* populations have been estimated to reach as high as 10 µM^[26–28]^, but it is possible that conditions at the point of cAMP release—especially at high cellular density—may produce much higher concentrations. We therefore opted to combine aptamer switches tuned to have different dynamic ranges within a single hydrogel, thereby enabling detection of both highly concentrated cAMP point-release within cell clusters and subsequent diffusion of the analyte. We functionalized our hydrogel with a 1:1 mixture of T9 and T33 aptamers and observed a cAMP-sensitive fluorescent response that combined the dynamic range of both the high-affinity and low-affinity receptors, achieving both an LOD of 7.9 µM and receptor saturation at >10 mM cAMP (**Figure 2A**, T9+T33). We used this T9+T33 aptamer system for subsequent experiments as it was best for achieving high sensitivity to changes in cAMP concentration across a >1,000-fold dynamic range (∼10 µM–10 mM).

### Characterization of the cAMP-responsive hydrogel

To ensure that the SEMAPHORE system was suitable for long-term continuous measurements, we investigated the stability of hydrogel conjugation and the response of aptamer switches in the hydrogel. Aptamer switches immobilized within the hydrogel exhibited stable confinement within the matrix over 12 h of immersion in buffer (**Figure S3a**). When we exposed these hydrogels to eight continuous cycles of cAMP-free buffer or buffer spiked with 10 mM cAMP, we observed reproducible fluorescent signaling across all cycles (**Figure S3B**). We then validated the spatiotemporal accuracy of SEMAPHORE by using a chemotaxis chamber to introduce a controlled concentration gradient between reservoirs containing 0 and 1 mM cAMP on either side of the hydrogel (**Figure S4**). We observed a steadily propagating wave of fluorescent signal increase that is consistent with cAMP diffusion through the hydrogel-filled chamber. The calculated diffusion coefficient (7.5 cm^2^/s) of 1 mM cAMP in SEMAPHORE was in the range of previously observed values for ATP in aqueous solution (3.5–7.5 cm^2^/s)^[29,30]^. Importantly, the high and uniform density of aptamer receptors within our hydrogel ensured that the spatial resolution of SEMAPHORE is only limited by the microscopy setup (see Methods for details).

We next compared the sensitivity of spatially resolved measurements across this cAMP gradient using SEMAPHORE. To establish ground truth for our cAMP gradient across the field of view, we measured the diffusion gradient established by free Alexa Fluor 488 dye prepared under the same conditions as our cAMP experiments. This offered a directly measurable surrogate small molecule with similar diffusive properties to cAMP. We then measured the cAMP gradient response for SEMAPHORE systems functionalized with differently tuned aptamers (T9, T33, and the 1:1 mixture of T9 and T33) and converted the fluorescence values measured across the field of view back into predicted cAMP concentrations based on the calibration curves obtained above for our aptamer switches. The T9/T33 hydrogel most accurately described the actual cAMP concentrations for the entire field of view as estimated from the Alexa Fluor 488 control experiment (**Figure 2B**). In contrast, single-aptamer measurements failed to do so across the full field of view due to their limited dynamic range. The T9 system lacked the sensitivity to detect low cAMP concentrations (0–60 µM), while the T33 system became saturated at high concentrations (60 μM–1 mM).

To measure the temporal resolution of our platform, we continuously captured fluorescence images from our T9/T33 hydrogel while varying cAMP concentrations over time (**Figure 2C**). We applied timed injections of 2.5, 5, and 10 µM cAMP onto the gel surface, with washing between each injection. Our system responded rapidly, reaching 90% of the steady-state fluorescence value within 20 s of cAMP addition. After washing away the cAMP, the signal was quenched to 10% of background within 30 s. In comparison, when the T33 aptamer switch was immobilized directly on the glass surface with no hydrogel to block the diffusion of cAMP to the receptor, the fluorescent switching response to 100 µM cAMP saturated within 3 s and upon washing, recovered to baseline within 9 s (**Figure S5**). These faster kinetics in the absence of hydrogel indicate that the response rate of SEMAPHORE is governed by the diffusion kinetics of cAMP through the hydrogel, rather than limitations imposed by the aptamer switch. The signal-on and -off kinetics observed in SEMAPHORE are therefore sufficient for real-time monitoring of *Dictyostelium* signaling, as previous observations have shown that the cAMP wave behavior exhibited by *Dictyostelium* cells has a typical period of 6 min/wave in the aggregation phase^[22,31]^.

### Imaging cAMP levels in *Dictyostelium* cells chemotaxing towards exogenously applied cAMP

Having demonstrated the spatiotemporal imaging performance of SEMAPHORE, we next introduced *Dictyostelium* cells into our system. *Dictyostelium* cells subjected to nutrient starvation directionally migrate in the direction of increasing cAMP concentration gradients^[20,21]^. We therefore used SEMAPHORE to analyze this chemotactic response while simultaneously tracking the spatiotemporal distribution of cAMP during measurement. *Dictyostelium* cells starved for 4 hrs to initiate their developmental program were transferred to the hydrogel, after which 10 µM cAMP was applied to a point on the hydrogel using a micropipette. We then observed the time-dependent behavior of the cells using the brightfield channel of an inverted fluorescence microscope, while also monitoring fluorescence to measure the cAMP gradient via SEMAPHORE (**Figure 3A–B**, cAMP applied outside the bottom-right corner of each image).

**Figure 3.**
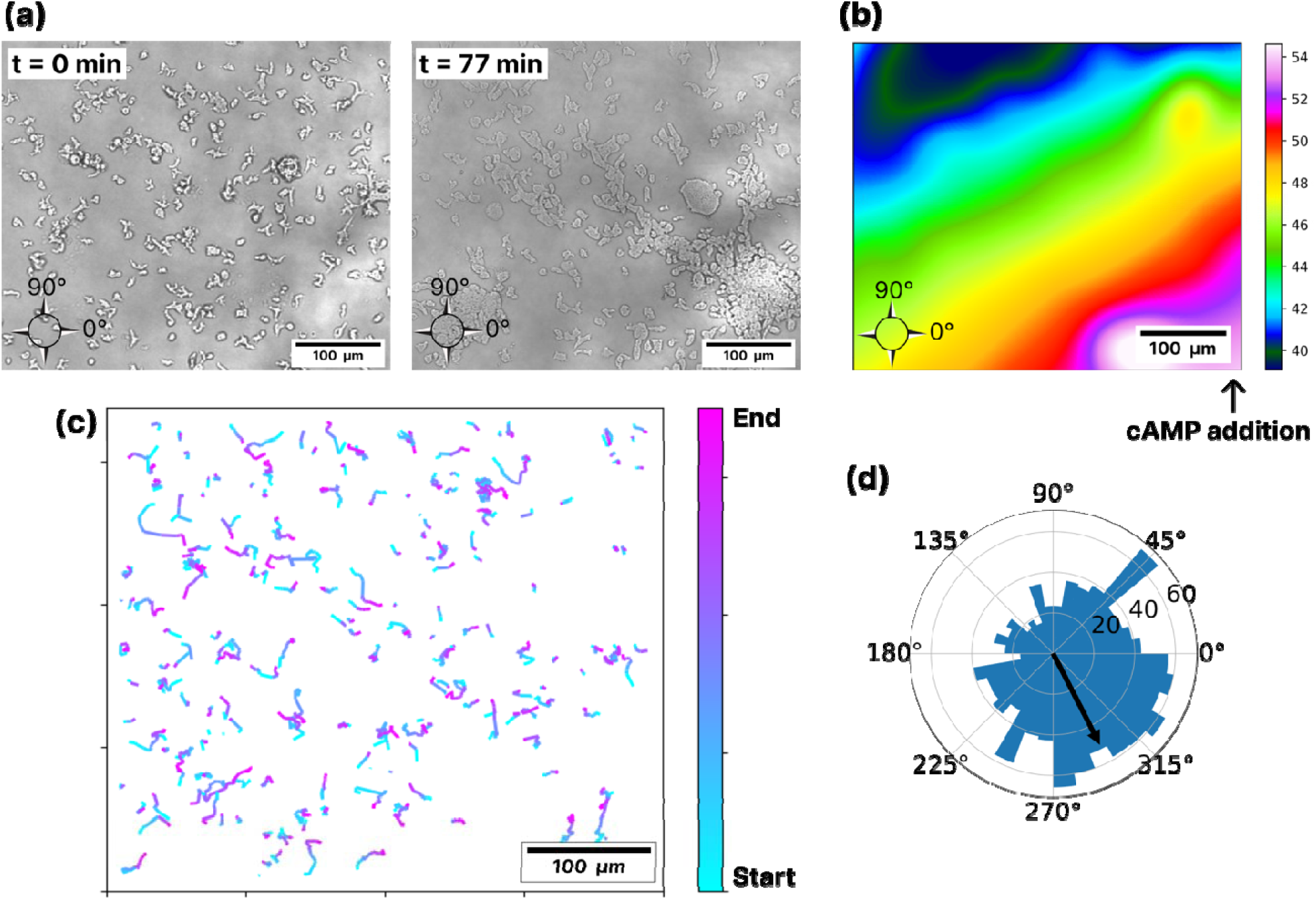
Monitoring *Dictyostelium* chemotaxis in response to exogenous cAMP using th SEMAPHORE system. (a) Brightfield images of *Dictyostelium* cells on the SEMAPHORE at 0 min (left) and 77 min (right) after injection of cAMP (complete brightfield recording provided as **Movie S1**) and (b) average fluorescence gradient (values correspond to measured fluoerscence intensity) measured by SEMAPHORE over the course of the 77 min measurement after applying cAMP (complete fluorescent recording provided as **Movie S2**). The cAMP injection was located just outside the lower-right corner of the hydrogel as indicated in panel (b). (c) Traces showing the migration of randomly selected 200 cells, colored by starting and ending positions (from cyan to pink) (d) Polar coordinate histogram (arbitrary units) of start-to-end cell angular movement (weighted by velocity). Arrow shows the direction of the average fluorescent gradient across the 77 min measurement calculated from the SEMAPHORE recording.

Our results confirmed the expected relationship between the direction of cellular migration and the exogenous cAMP gradient. We used automated analysis of the brightfield data (see Supporting Methods) to track the migration of individual cells over time as the population progressed from unicellular cells into large aggregates over the course of 77 min (**Fig 3A, C, Movie S1)**. Through direct measurement of the SEMAPHORE fluorescent response, we determined that the average angle of the cAMP gradient within the field of view was 297° (**Fig 3B, Movie S2**). After plotting directional cell movement on a polar histogram, weighted by velocity, we observed a clear bias in cell motion toward directions ranging between 225° and 50° (**Figure 3D**). This distribution matches well with the average measured cAMP gradient angle (black arrow in **Fig 3D**), illustrating the expected cell migration towards higher cAMP concentrations. In contrast, cells moved randomly when exposed to a uniform distribution of applied cAMP (no gradient across a majority of the field of view) for a similar duration (**Figure S6, Movies S3, 4**). The measured gradients may deviate from the exact applied cAMP gradients due to interference from both endogenous cAMP secretions from cells, as well as cAMP degradation from secreted adenylyl cyclases which degrade cAMP and may produce some cross-reactive adenosine derivative species. However, the agreement between SEMAPHORE and brightfield cell movement measurements demonstrate clearly that we can simultaneously measure both cell behavior and the extracellular molecular signals driving that behavior using SEMAPHORE.

### Monitoring extracellular cAMP signals underlying chemotaxis in real time

Finally, we used SEMAPHORE to monitor *Dictyostelium* behaviors driven by endogenous rather than exogenous cAMP signals. Under starvation conditions, individual *Dictyostelium* cells respond to and secrete cAMP, giving rise to characteristic cAMP waves and migration of individual cells to assemble into a multicellular organism. This process unfolds over the course of ∼24 h, with individual cells forming characteristics streams that come together into tight aggregates within 3–7 h after the onset of starvation^[22,23]^. The cAMP waves underlying aggregation have previously been studied indirectly by monitoring either cell positions, waves at a single point in time through isotope dilution-fluorography, or intracellular cAMP levels^[22,31,32]^. We hypothesized that SEMAPHORE enables direct recording of the extracellular cAMP signaling dynamics that drive streaming and aggregation.

We starved *Dictyostelium* cells in nutrient-free buffer for 2 hrs and transferred them to a nutrient- and cAMP-free SEMAPHORE hydrogel for observation. Over the course of 7 hrs, we collected brightfield images every 10 secs to observe the behavior of cells (**Movie S5)**. Simultaneously, we captured fluorescent images from the SEMAPHORE platform to measure extracellular cAMP concentrations (**Movie S6)**. Brightfield images showed chemotactic behaviors consistent with results from the literature (**Fig. 4A, Top**)^[23,31]^. Between 2–5 hrs after the beginning of starvation, we observed predominantly single cell dynamics with some preliminary aggregation. At 5–7.5 h post-starvation, we observed active migration and streaming into aggregates, giving way to mounds and tipped mounds indicating the onset of the slug phase by 7.5–9 h post-starvation.

**Figure 4.**
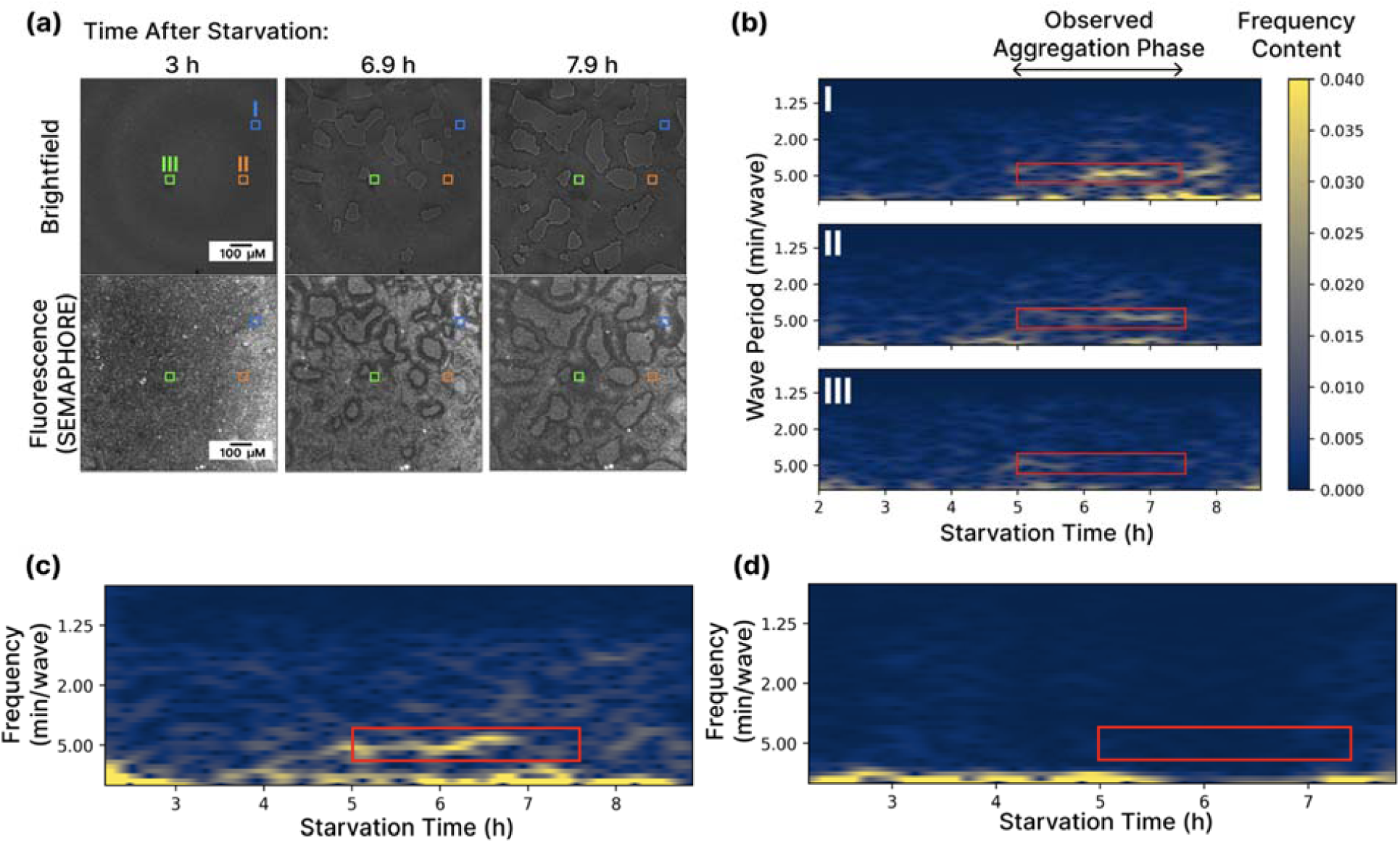
Imaging extracellular cAMP signaling during *Dictyostelium* aggregation in real-time. (A) Brightfield (top) and fluorescence images (bottom) of cells migrating on the SEMAPHORE hydrogel at 3 h (top), 6.9 h (middle), and 7.9 h (bottom) after the onset of starvation. (B) Heat-map of the cAMP signal frequency content across time for the highlighted regions of panel A. Red inset boxes correspond to the times of observed aggregation, and the previously reported frequency ranges of cAMP oscillation during *Dictyostelium* chemotaxis. (C) Map of Fourier-transformed frequencies for the Moran’s I statistic, which quantifies the oscillation behavior of the entire population over time, unbiased by the selection of specific regions within the image. (D) Moran’s I analysis for a control SEMAPHORE experiment in which we used a scrambled aptamer switch with no cAMP affinity. Red rectangles in panels B-D highlight the time periods over which aggregation was observed as well as the observed oscillation frequencies.

SEMAPHORE allowed us to augment these observations with direct recordings of the extracellular cAMP waves underlying migration and aggregation **(Movie S6)**. We selected regions of interest corresponding to either aggregation centers (**Figure 4A**, regions I & II) or regions that cells migrate away from over the course of imaging (**Figure 4A**, region III). We analyzed local cAMP wave behavior by averaging the fluorescence within these regions and applying a short-time Fourier transform (STFT) to quantify the frequency of cAMP oscillations across the duration of chemotaxis (**Figure 4B**, see Methods). The most prominent frequencies are marked by the yellow regions in **Figure 4B** and correspond to the characteristic cAMP oscillations that drive *Dictyostelium* aggregation. At both aggregation centers, we observed high-amplitude cAMP oscillations with a period of 2.8–8.3 min/wave during the phase of most rapid aggregation (5–7.5 h post-starvation). Both the timing of onset for oscillations and the periodicity observed here are consistent with previous observations obtained with established techniques for observing cAMP response, which typically calculate a period of 6 min/wave^[22,31]^. These high-frequency oscillations have the largest amplitude during the periods of fastest aggregation (red inset boxes in **Figure 4**). All regions exhibited slow cAMP fluctuations (>10 min/wave, < 0.1 s^-1^) during early starvation (<5 hours) and post-aggregation slug phases (>8 hours) owing to a lack of coordinated migration. We also observed minimal oscillatory behavior far from the aggregation centers, as noted by the weak frequency content in region III. It is possible that cAMP degradataion by secreted adenylyl cyclases may lead to the slow accumulation of other adenosine derivatives for which the aptamer has cross-reactive binding. However, the close agreement between our most prominent measured frequencies and known cAMP oscillation frequencies from previous literature indicates that if these effects occur, they are minor and do not interfere with our ability to observe the relevant biological behavior.

As a control for the specificity of SEMAPHORE in terms of cAMP detection, we also performed these experiments using a non-cAMP-responsive hydrogel incorporating DNA receptors with a matching structure but with scrambled cAMP-recognition aptamer sequences. We observed the same multi-stage aggregation behavior in our brightfield images (**Figure S7A, Movie S7**) indicating that the aptamer sequence has minimal effect on the chemotactic behavior. However, we observed only low-level background fluorescence in our SEMAPHORE measurements (**Figure S7B, Movie S8**), confirming that the oscillations described above were specifically associated with aptamer switch binding of cAMP. Further, before the onset of starvation-driven aggregation (hours 2-4 post-starvation in **Figure 4**) we observe no characteristic fluorescence oscillations, confirming that the cAMP readout generated by our SEMAPHORE measurements is directly correlated with *Dictyostelium* migration.

To eliminate the possibility of bias arising from the selection of specific aggregation regions for frequency analysis, we also performed global characterization of cAMP oscillation throughout the full *Dictyostelium* population. To do this, we calculated the Moran’s I statistic, which distills the spatial distribution of cAMP across the entire field of view into a single value at each point in time^[33,34]^. This value quantifies the spatial autocorrelation of cAMP: how closely cAMP values at points across the image correlate with the cAMP value of their close neighbors. A high Moran’s I value (close to 1) indicates locally synchronized secretion of cAMP across the population, while a value of 0 indicates no spatial coordination. We calculated the Moran’s I value across the full time-course (2–9 h post-starvation) and then again applied the STFT method to these Moran’s I values to study the frequency content of global cAMP oscillation. As with the localized measurements, we observed the strongest high-frequency oscillations during the times of fastest aggregation (4.8–7 h; **Figure 4C**). These oscillations had a period of 4–6 min/wave, in strong agreement with both our local measurements and previous findings^[22,23,31]^. No characteristic oscillations were observed when this global analysis was applied to data collected using a scrambled aptamer sequence (**Figure 4D**). This suggests that our SEMAPHORE platform is specific for the detection of cAMP waves and is not confounded by other nonspecific effects.

## Conclusion

SEMAPHORE provides a sensitive method for generating continuous ‘molecular movies’ of extracellular communication. Our platform combines fluorescent aptamer-based molecular switches with a biocompatible hydrogel matrix to achieve continuous, multi-hour recordings of signaling molecule secretion by cells over an entire 2D field of view using fluorescence microscopy. By combining a pair of aptamer switches with distinctively tuned cAMP affinity, we were able to achieve sensitive measurement across a broad dynamic range, allowing us to monitor a wide range of cAMP concentrations applied to or generated by *Dictyostelium* cell populations. We observed close agreement of our SEMAPHORE measurements with current experimental and theoretical knowledge of *Dictyostelium* chemotaxis under conditions of both exogenous and endogenous cAMP signaling. We used our system to directly record the extracellular cAMP signals driving the chemotactic behavior of cells under starvation conditions, while also recording cell movement using brightfield imaging. cAMP-driven chemotaxis has been extensively studied using brightfield cell recordings, intracellular cAMP measurements, and single-timepoint extracellular cAMP measurements^[22,23,27,31,35]^, but our system now makes it possible to continuously record the extracellular cAMP waves that drive this collective cell behavior while also observing the chemotaxis process itself. These direct extracellular observations have the potential to add a new dimension to studies of cellular function.

This approach should be generalizable for the monitoring of other extracellular signaling molecules for which an aptamer switch is either available or can be generated. Nevertheless, several challenges will need to be addressed when extending this platform to observe new biological phenomena. Although the hydrogel matrix can readily incorporate new fluorescent aptamer switches with specificity for diverse molecular signals, careful optimization of aptamer concentration, sensitivity, and kinetic response will be required for each new switch design. When applying SEMAPHORE to a new biological system, properties such as the analyte dynamic range or the timescale of secretion may be poorly understood or even fully unknown. This means that the aptamer response must exhibit a rapid response and high sensitivity across a broad concentration range to ensure that the platform can detect the full range of possible signaling activity. This could pose a fundamental challenge, as sensitive, high-affinity receptors typically exhibit slow binding kinetics. Fortunately, there is recent progress towards developing aptamer tuning methods that can decouple a receptor’s sensitivity from its kinetic response^[17]^, as well as algorithmic strategies that enable accurate determination of rapidly-changing concentrations even from the out-of-equilibrium dynamics of ‘slow’ receptors^[36]^. Overcoming these various challenges would make SEMAPHORE a broadly useful tool for studying extracellular signals in a wide range of important biological phenomena ranging from cell fate determination^[37,38]^ to the collective behavior of cellular populations^[39]^ to processes such as neurotransmission and immune cell communication^[7,40]^.

## Methods

### Materials

HPLC-purified oligonucleotides modified with methacrylamide, fluorophore, and quencher were purchased from Integrated DNA Technologies. All oligonucleotides were resuspended in nuclease-free water and stored at −20 °C. All sequences used in this work are shown in **Table S1**. Unless otherwise specified, all other chemicals were purchased from Thermo Fisher Scientific. All cell experiments were conducted in development buffer (DB): 10 mM Tris-HCl (pH 6.5), 2 mM MgCl_2_, 1 mM CaCl_2_. *D. discoideum* (strain-DBS0235747, axenic) was purchased from Dicty Stock Center. LoFlo Medium powder was purchased from ForMedium. All images for aptamer switch testing in the absence of cells were recorded by an Olympus IXplore standard inverted fluorescent microscope with an Andor sCMOS camera. We used Leica Dmi-8 for the continuous *in situ* imaging of brightfield and fluorescence images. All experiments were performed in triplicate unless otherwise noted.

### Fabrication of anchor DNA-conjugated hydrogels

Prior to polymerization, the inhibitor (<100 ppm 4-methoxyphenol) of PEGDA (M_n_ 700 g mol^-1^) was removed by passing through a column with neutral alumina. 0.3% (w/w) 2-hydroxy-4-(2-hydroxyethoxy)-2-methylpropiophenone (98%), 7% (w/w) PEGDA, and 0.5 µM oligonucleotides (functionalized with a methacryl group at the 5’ end and BHQ2 at the 3’ end) were mixed in PBS pH 7.4 buffer to yield the pregel solution immediately before use. The mixture was purged with nitrogen gas for 3 min. The optic grade glass was immersed in piranha solution (3:1 sulfuric acid:hydrogen peroxide) for 30 min, and then rinsed with DI water and acetone. After drying the glass with a nitrogen gas blower, the glass was immersed in 1% (v/v) 3-(trimethoxysilyl)propyl methacrylate in acetone for 30 min. The methacrylate-functionalized glass was rinsed with acetone, dried via filtered dry nitrogen gas, and finally stored in the -20℃ freezer before we use. Two 50-µm-thick strips of Kapton tape were attached to one surface of a clean glass slide, spaced 1cm apart. The methacrylate-functionalized glass was then placed atop this assembly, leaving a 50-µm gap with an area of 20 x 20 mm. The pregel solution was introduced into this gap between the sandwiched glass slides with a pipette via capillary force, after which the solution was exposed to 365 nm irradiation for 2.5 min using a UV hand lamp. After peeling off the clean glass slide, the hydrogel remained covalently bonded to the methacrylate-treated glass. The gel-coated slide was rinsed three times with PBS (pH 7) to remove any residues, and then immersed in PBS and in a 100% humidity closed chamber at 4 °C until use. This process is shown in **Figure S8**.

### Dictyostelium growth

For the imaging experiment, frozen cell stocks were defrosted in a 37 °C water bath only until the stock was partially melted to allow removal from the tube without heating the cells above room temperature. The cells were then gently redispersed in LoFlo medium. The cells were then grown to a density of ∼5 x 10^6^ cells/ml at 22–25 °C prior to each experiments.

### Assembly and preparation of the SEMAPHORE imaging chamber

A 4-mm-thick film of polydimethylsiloxane (PDMS; Sylgard 184, Dow Corning) was cut into 2 x 2 cm pieces. A central hole was drilled using a punch biopsy tool with a 6 mm diameter (Integra™ Miltex™ Standard). The PDMS film was then positioned atop the hydrogel, with the puncture positioned at the center of the hydrogel to serve as an imaging window. The PDMS film was then affixed to the slide with epoxy glue applied to its borders. 100 µl of 0.5 µM aptamer switch was pipetted onto the imaging window and then incubated for 3 h under moist conditions by storing in a closed Petri dish also containing wet wipes. The imaging window was then rinsed with PBS, filled with 80-100 µl of PBS and aged for 3 h, and then rinsed again with PBS. This process is shown in **Figure S8**.

### Microscopy protocol for simultaneous brightfield and fluorescent imaging

For *in situ* brightfield and fluorescence imaging, we used an inverted light microscope (Leica Dmi-8) with a 10× objective lens (HC PL APO CS2 10x/0.40 DRY, numerical aperture 0.4), and controlled using LAS X Life Science Microscope Software. Samples were excited with a 522-nm laser (OPSL552, intensity 5%). The emission signal was separated using a dichroic beam splitter (DD 488/552). For the photomultiplier tube (PMT) transmission and detector channels, we used 560– 725 nm bandwidth (pinhole = 53.1 µm; pinhole Airy = 1 AU; emission wavelength for pinhole Airy = 580 nm). We captured 8-bit 1024 × 1024 pixel images (pixel size = 1.136 µm) with a scan speed of 400 Hz, 84 total exposures (2 channels, 42 frames), and a multiplication gain of 900–1100.

### Fluorophore and quencher degradation test

We prepared all samples on the 96 well microplate (Corning® 96-well Half Area Black Flat Bottom Polystyrene Microplate). The samples are adjusted to the total 50 µl into each wells. For the test of Cy3 dye, 200 µM of T33 in PBS buffer with photoinitiator (+PI), 0.3% (w/w) 2-hydroxy-4-(2-hydroxyethoxy)-2-methylpropiophenone, and without (-PI) is used. The BHQ-2 sample is same except the T33 is switched to the anchor strand. After covering the well using the plastic coverslip (VWR^TM^), the samples to be UV irradiated are UV exposed for a period of time using a UV hand lamp located directly above the sample plate. Then, before analyzing the BHQ-2 sample, the same molar amount of T33 is added, adjusted to the 55 µl in total, and stabilized for 10 minutes. The fluorescence intensity for all samples were measured at 25D°C on a Synergy H1 microplate reader (BioTeK). Emission spectra were monitored in the 550–700 nm range with Cy3 excitation at 530 nm and a gain of 100.

### Long term stability test

The SEMAPHORE samples are same except that the PDMS cover is not used. The physically trapped aptamer/hydrogel was fabricated by following the same recipe of SEMAPHORE except that there is no acrydite functional group at the 5’ end of anchor strand. The pristine hydrogel was prepared by polymerization of 0.3% (w/w) 2-hydroxy-4-(2-hydroxyethoxy)-2-methylpropiophenone, 7% (v/v) PEGDA in PBS pH 7.4 buffer. The samples on the surface-treated glass are immersed in PBS buffer in Petri dishes, and then incubated during a period of time (12h per one cycle) under dark condition on an orbital shaker at 30-60 rpm. Images were taken by an Olympus IXplore standard inverted fluorescent microscope with an Andor sCMOS camera (4x objective lens, excitation power = 3, Nikon Cy3 bandwidth filter). We captured 16-bit 1024 × 1024 pixels images with an exposure time of 500 ms and 2x2 binning. Intensities was measured by median values from the resulting grayscale images. For the multiple cycling test of SEMAPHORE, the solution buffer was switched from 0 to 10mM of cAMP and vice versa after intensity measurement for each cycles.

### ISD switch binding affinity measurements

For these experiments, the SEMAPHORE system was assembled in a sticky-slide chemotaxis chamber (ibidi). After removing the protective film, a methacrylate-modified coverslip was attached to the adhesive surface. Then, approximately 30 µL of pregel solution was injected into the central observation chamber (2 mm x 1 mm x 70 µm) using micro pipette, and then exposed to UV (365 nm) for 2.5 min. The two reservoir chambers (130 µl each) were filled with DB and kept at room temperature for 3 h. After removing the solution, the reservoirs were refilled with 130 µL of 0.5 µM cAMP-responsive aptamer switches dissolved in PBS and incubated for 24 h at 4 °C. The reservoirs were then filled with DB solution, stored for 1 h at room temperature, after which the solution was removed via pipette. This was repeated three times, and the DB solution was kept in the reservoirs overnight for the last wash. For the experiment, one reservoir was filled with DB containing cAMP at the desired concentration, whereas another reservoir was filled with DB only. Images were captured by an Olympus IXplore standard inverted fluorescent microscope with an Andor sCMOS camera (4x objective lens, excitation power = 3, Nikon Cy3 bandwidth filter). We captured 16-bit 1024 × 1024 pixels images with an exposure time of 200 ms. We observed a 1 mm × 2 mm central field of view.

### Analysis of binding properties

Raw fluorescence (I) vs. [cAMP] data for each aptamer switch construct were fitted to a Langmuir isotherm binding model to extract the binding affinity of the construct:

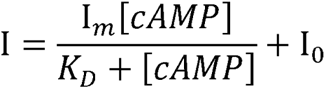

where I_0_ is the background signal at [cAMP] = 0, I*_m_* is the maximum signal at saturating target concentration, and *K_D_* is the dissociation constant. We corrected for small variations in aptamer concentration between the samples by normalizing all binding curves to the target-free background fluorescence. These normalized binding curves were produced by correcting all data to 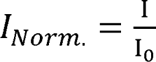, followed by replotting with a Langmuir isotherm fit that was normalized by the same factor I_0_.

For the mixed aptamer system, where the two different aptamers are described by K*_D_*_,1_ and K*_D_*_,2_, we fitted cAMP-dependent curves with the functional form:

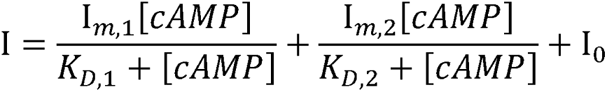

### Aptamer functionalization of glass surfaces for kinetic tests in the absence of hydrogel

To prepare the substrate for aptamer switch immobilization, a glass cover slip was treated with 3:1 mixture of 98% sulfuric acid and hydrogen peroxide for 1 h, rinsed with DI water for 3 times and dried by filtered N_2_ gas. Then, the substrate was treated with 1% VECTABOND reagent (Novus Biologicals) in acetone for 5 minutes, rinsed with DI water 3 times and dried by filtered N_2_. 100 µL solution containing 0.2 mg biotin-PEG-SVA-5000 (Laysan Bio), 10 mg PEG-SVA-5000 (Laysan Bio) and 0.825% sodium bicarbonate was sandwiched by two pieces of VECTABOND treated cover slips for 3 hrs. The slip was rinsed with DI water 3 times dried by air and stored at - 20 °C until used. Next, PEG-modified glass cover slip was assembled with adhesive chamber (GBL612107, HybriWell adhesive chamber). Through the inlet of the chamber, 0.2 mg/mL NeutrAvidin solution (Thermo Scientific) was injected and incubated for 5 minutes, then rinsed with fresh DB 3 times. 100 nM of anchor DNA sequence with biotin 5’ end group (5’/biotin/TTTTTGCTTCGGCTCGTATA/BHQ2/3’) was injected and incubated for 5 minutes, rinsed with fresh DB. 100 nM aptamer switch sequence (T33) was injected and incubated for 5 minutes, rinsed with fresh DB. The system was kept in DB at 4 °C until before we use.

### Temporal resolution analysis

Using either the SEMAPHORE assembly or the directly aptamer-modified glass within the PDMS chamber, we applied 50 µl of DB to the imaging window and captured the fluorescence images to obtain the blank values. I_0_ was calculated by averaging the three different median values from distinct images of the sample with fresh DB from the resulting grayscale images. The desired cAMP concentration was achieved by adding 10 µl concentrated solution in DB to 50 µl of DB in the imaging window. To refresh the DB in the tests using the SEMAPHORE platform, imaging was paused and the solution was changed to 50 µl of DB. After 10 s, imaging was resumed. Imaging was not paused during the aptamer-on-glass measurements to allow the rapid kinetics to be captured. Images were acquired every 1 s (except during the refreshing period) using an Olympus IXplore standard inverted fluorescent microscope with an Andor sCMOS camera (4 x objective lens, excitation power = 3, Nikon Cy3 bandwidth filter, 16-bit 1024 × 1024 pixels, binning 2x2, and an exposure time of 200 ms). Intensity values were calculated using a median values of grayscale image of each frame.

### Imaging during exogenous cAMP addition and chemotaxis

Cells were detached from the dish surface by gentle pipetting and then washed with DB once by centrifugation for 3 min at 200–300 rcf. The cells were resuspended in fresh DB (∼2 x 10^7^ cells/ml) and given pulses of 30 nM cAMP in DB at 6 min intervals for 5 hrs^[22]^. The cells were then incubated for 30 min, after which we replaced the media with fresh DB at room temperature. After removing some of the cell aggregates by pipetting gently, 70 µl of cell-dispersed DB solution was applied to the SEMAPHORE imaging chamber. The cells were then incubated for 10 min at room temperature to facilitate attachment to the surface. The solution was carefully removed via micropipette, and any remaining liquid was absorbed using cleanroom wipes. Subsequently, 0.5 µL of 10 µM cAMP was added to one corner of the field of view (1162.5 x 1162.5 µm) using a WPI micropipette tip with a diameter of 10 µm. Then, the exposed surface of the imaging window was covered with an optical-grade cover slip (ibidi) to retain moisture during the measurement. Images were captured at the location where the cAMP-driven fluorescence gradient was created by changing the position of the field of view.

### Imaging during endogenous cAMP-driven chemotaxis

Cells were carefully detached from the surface of the dish by gentle pipetting and washed three times with DB by centrifuging for 3 min at 200–300 rcf. The cells were then starved for 2 h in DB at approximately 2 x 10^7^ cells/ml at room temperature. After gently pipetting to remove cell aggregates, 70 µl of the cell-dispersed DB solution was applied to the imaging window and incubated for 10-15 min to allow for attachment to the surface at room temperature. The solution was removed via micropipette, and any residual liquid was soaked up using cleanroom wipes. Then, the imaging window was covered with an optical-grade cover slip. The entire system was placed in a closed Petri dish containing wet wipes to keep it hydrated during measurements. Simultaneous brightfield and fluorescence imaging were acquired for approximately 7 hrs using an inverted light microscope.

### Automated cell tracking

We first masked our brightfield images using Cellpose Prediction for 2D v0.2 with the following parameters: model: Nuclei, omniflag: True, diameter: 10, flow threshold: 1, cell probability threshold: -5. The binary masks were imported as a 16-bit image stack into Fiji. Trackmate was then used to track cellular movement. First, the Difference of Gaussian (DoG) detector was used with the following parameters: estimated object diameter: 12, quality threshold: 8, sub-pixel localization: True. Subsequently, the Sample Linear Assignment Problem (LAP) tracker was used with the following parameters: linking max distance: 5, gap-closing max distance: 10, gap-closing max frame gap: 2. For experiments performed in the absence of exogenous cAMP, the larger image frames meant that we had to instead run Cellpose Prediction with the following parameters: model: Nuclei, omniflag: True, diameter: 4, flow threshold: 1.1, cell probability threshold: -6. DoG detector was used with the following parameters: estimated object diameter: 2, quality threshold: 3, sub-pixel localization: True. Sample LAP tracker was used with the following parameters: linking max distance: 15, gap-closing max distance: 15, gap-closing max frame gap: 2.

Finally, the tracks were exported subjected to analysis and plotting. First, cells that were detected within 10 pixels of the image boundaries were removed from tracking as they may have left the imaging region. The velocity of each tracked cell was then calculated based on the Euclidean distance traveled divided by the number of frames passed since the last tracked frame:

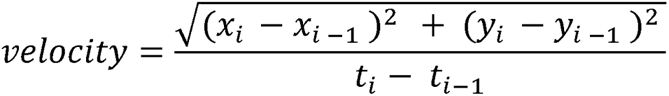

where x_i_, y_i_ is the pixel location of a cell at frame t _i_, and x _i–1_, y _i–1_ is the pixel location of a cell at frame t _i–1_. The angle of each tracked cell was calculated by the arctan of the current location and previous tracked location. Histograms were weighted by velocity and plotted on polar coordinates as Windrose plots.

### Moran statistics

The Moran’s I statistic was calculated using the python library pysal by:

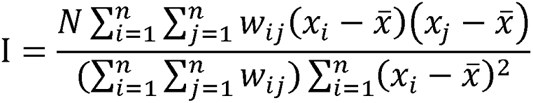

where *x_i_* and *x_j_* are the fluorescence intensity values, 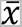 is the mean fluorescence intensity, w is the spatial weight matrix defined by rook spatial proximity, and N is the number of observations/pixels.

### Fluorescence image processing

In this section, we describe the data processing done to estimate the gradients of the images in Fig. 3b and Fig. S6c. This image processing was necessary because the gradient values were very small compared to the noise levels. If one were to naïvely estimate the gradient as the difference in RFU of two neighboring pixels, the noise of the two pixels would be additive, which would overwhelm the signal of approximately 0.05 RFU/px. We assume two things for our data processing: First, the true image gradients are ‘shallow’, meaning that ‘steep’ high-frequency components of the image are mostly noise. Second, there is little photobleaching across time in the underlying experiments, so variability across time is mostly noise. To remove the noise from the high-frequency components, we apply the scipy function gaussian_filter to each frame individually^[41]^. This acts as a low-pass filter. We chose a sigma of 22.75µm, which corresponds to 20px for Fig 3b, and 10px for Fig S6c. To remove the noise from frame-to-frame variability, for every pixel, we calculate the average pixel value across time, yielding a single image. Using this denoised, smoothed image, we then estimate the gradient values using simple first order finite differences. To finally display this image, we first crop the image to the region of interest (for Fig. 3b this is a sub-region, for Fig. S6c this is the entire image), and then choose the lower and upper colormap limits to be equal to the 1^st^ and 99^th^ percentile of the pixel intensities, respectively.

### Short-time Fourier transform (STFT)

We use the STFT to resolve which frequencies are present in a signal over time. STFT entails the calculation of multiple Fourier transforms on subsections of the signal. While the reduction in analyzed signal length reduces the resolution of the Fourier transform, the STFT allows us to gain insight into when a given frequency is present in a signal. In this work, the STFT was performed using Scipy with a window size of 256 samples with an overlap of 255^[41]^.

## Acknowledgements

We thank Professor Carole A. Parent for providing guidance on the biology of *Dictyostelium discoideum* cells and microscopy-based functional cell assays. HTS gratefully acknowledges financial supported by Helmsley Charitable Trust and the Wellcome LEAP SAVE program. This work was also supported by the National Research Foundation of Korea(NRF) grant funded by the Korea government(MSIT) (No. RS-2023-00251670). I.A.P.T. was supported by the Medtronic Foundation Stanford Graduate Fellowship and the Natural Sciences and Engineering Research Council of Canada (NSERC, 416353855). S.S.N. acknowledges support from SGF (Stanford Graduate Fellowship in Science & Engineering) and the NSF Graduate Research Fellowship Program (GRFP).

## References

[1] B. N. G. Giepmans, S. R. Adams, M. H. Ellisman, R. Y. Tsien, Science 2006, 312, 217.

[2] J. R. James, R. D. Vale, Nature 2012, 487, 64.

[3] C. Uttamapinant, J. D. Howe, K. Lang, V. Beránek, L. Davis, M. Mahesh, N. P. Barry, J. W. Chin, J Am Chem Soc 2015, 137, 4602.

[4] J. M. Baskin, J. A. Prescher, S. T. Laughlin, N. J. Agard, P. V. Chang, I. A. Miller, A. Lo, J. A. Codelli, C. R. Bertozzi, Proceedings of the National Academy of Sciences 2007, 104, 16793.

[5] E. M. Sletten, C. R. Bertozzi, Angew Chem Int Ed Engl 2009, 48, 6974.

[6] J. S. Dickschat, Nat. Prod. Rep. 2010, 27, 343.

[7] S. E. Hyman, Curr Biol 2005, 15, R154.

[8] E. Potma, W. P. de Boeij, P. J. van Haastert, D. A. Wiersma, Proc Natl Acad Sci U S A 2001, 98, 1577.

[9] M. P. McDonald, A. Gemeinhardt, K. König, M. Piliarik, S. Schaffer, S. Völkl, M. Aigner, A. Mackensen, V. Sandoghdar, Nano Lett 2018, 18, 513.

[10] M. P. Raphael, J. A. Christodoulides, J. B. Delehanty, J. P. Long, J. M. Byers, Biophys J 2013, 105, 602.

[11] J. Juan-Colás, I. S. Hitchcock, M. Coles, S. Johnson, T. F. Krauss, Proc Natl Acad Sci U S A 2018, 115, 13204.

[12] X. Li, M. Soler, C. Szydzik, K. Khoshmanesh, J. Schmidt, G. Coukos, A. Mitchell, H. Altug, Small 2018, 14, e1800698.

[13] Y. Shirasaki, M. Yamagishi, N. Suzuki, K. Izawa, A. Nakahara, J. Mizuno, S. Shoji, T. Heike, Y. Harada, R. Nishikomori, O. Ohara, Sci Rep 2014, 4, 4736.

[14] T.-G. Cha, B. A. Baker, M. D. Sauffer, J. Salgado, D. Jaroch, J. L. Rickus, D. M. Porterfield, J. H. Choi, ACS Nano 2011, 5, 4236.

[15] M. P. Landry, H. Ando, A. Y. Chen, J. Cao, V. I. Kottadiel, L. Chio, D. Yang, J. Dong, T. K. Lu, M. S. Strano, Nat Nanotechnol 2017, 12, 368.

[16] A. E. Rangel, A. A. Hariri, M. Eisenstein, H. T. Soh, Adv Mater 2020, 32, e2003704.

[17] B. D. Wilson, A. A. Hariri, I. A. P. Thompson, M. Eisenstein, H. T. Soh, Nature communications 2019, 10, 5079.

[18] Z. Tang, P. Mallikaratchy, R. Yang, Y. Kim, Z. Zhu, H. Wang, W. Tan, Journal of the American Chemical Society 2008, 130, 11268.

[19] A. M. Yoshikawa, L. Wan, L. Zheng, M. Eisenstein, H. T. Soh, Proc Natl Acad Sci U S A 2022, 119, e2119945119.

[20] J. M. Nichols, D. Veltman, R. R. Kay, Curr Opin Cell Biol 2015, 36, 7.

[21] C. A. Parent, P. N. Devreotes, Annu Rev Biochem 1996, 65, 411.

[22] K. Tomchik, P. Devreotes, Science 1981, 212, 443.

[23] T. Gregor, K. Fujimoto, N. Masaki, S. Sawai, Science 2010, 328, 1021.

[24] D. E. Huizenga, J. W. Szostak, Biochemistry 1995, 34, 656.

[25] F. Ricci, A. Vallée-Bélisle, A. J. Simon, A. Porchetta, K. W. Plaxco, Acc. Chem. Res. 2016, 49, 1884.

[26] M. Brenner, Dev Biol 1978, 64, 210.

[27] P. N. Devreotes, M. J. Potel, S. A. MacKay, Dev Biol 1983, 96, 405.

[28] G. Gerisch, U. Wick, Biochem Biophys Res Commun 1975, 65, 364.

[29] M. J. Hubley, R. C. Rosanske, T. S. Moerland, NMR Biomed 1995, 8, 72.

[30] W. J. Bowen, H. L. Martin, Arch Biochem Biophys 1964, 107, 30.

[31] G. Singer, T. Araki, C. J. Weijer, Commun Biol 2019, 2, 139.

[32] P. Fey, A. S. Kowal, P. Gaudet, K. E. Pilcher, R. L. Chisholm, Nat Protoc 2007, 2, 1307.

[33] P. a. P. Moran, Biometrika 1950, 37, 17.

[34] S. Sun, J. Zhu, X. Zhou, Nat Methods 2020, 17, 193.

[35] P. Devreotes, Science 1989, 245, 1054.

[36] N. Maganzini, I. Thompson, B. Wilson, H. T. Soh, Nat Commun 2022, 13, 7072.

[37] J. Oyler-Yaniv, A. Oyler-Yaniv, E. Maltz, R. Wollman, Nat Commun 2021, 12, 2992.

[38] C. Hetz, Nat Rev Mol Cell Biol 2012, 13, 89.

[39] A. Nakajima, S. Ishihara, D. Imoto, S. Sawai, Nat Commun 2014, 5, 5367.

[40] A. Trautmann, Sci Signal 2009, 2, pe6.

[41] P. Virtanen, R. Gommers, T. E. Oliphant, M. Haberland, T. Reddy, D. Cournapeau, E. Burovski, P. Peterson, W. Weckesser, J. Bright, S. J. van der Walt, M. Brett, J. Wilson, K. J. Millman, N. Mayorov, A. R. J. Nelson, E. Jones, R. Kern, E. Larson, C. J. Carey, İ. Polat, Y. Feng, E. W. Moore, J. VanderPlas, D. Laxalde, J. Perktold, R. Cimrman, I. Henriksen, E. A. Quintero, C. R. Harris, A. M. Archibald, A. H. Ribeiro, F. Pedregosa, P. van Mulbregt, Nat Methods 2020, 17, 261.

