## Supplementary Table and Figures for "Real-Time Spatiotemporal Measurement of Extracellular Signaling Molecules Using an Aptamer Switch-Conjugated Hydrogel Matrix"

**Contents**

**Table S1 |** Sequences of all constructs used in this work.

**Supplementary Figures S1–8**

**Supplementary Movies**

**Movie S1.** Brightfield image stack of stimulated dicty cells on SEMAPHORE for 77 min after generating an exogenous cAMP gradient (232 frames, frame interval 20 s, fps 20).

**Movie S2.** Median filtered fluorescence image stack of SEMAPHORE for 77 min after generating an exogenous cAMP gradient (232 frames, frame interval 20 s, fps 20).

**Movie S3.** Brightfield image stack of stimulated dicty cells on the SEMAPHORE hydrogel for 41 min after exogenous cAMP addition without a concentration gradient (248 frames, frame interval 10 s, fps 15).

**Movie S4.** Median filtered fluorescence image stack of the SEMAPHORE hydrogel for 41 min after exogenous cAMP addition without a concentration gradient (248 frames, frame interval 10 s, fps 15).

**Movie S5.** Brightfield image stack of 2 h starved dicty cells on the SEMAPHORE hydrogel under nutrient-free conditions for 6.9 h (2,471 frames, frame interval 10 s, fps 15).

**Movie S6.** Median filtered fluorescence image stack of 2 h starved dicty cells on the SEMAPHORE hydrogel under nutrient-free conditions for 6.9 h (2,471 frames, frame interval 10 s, fps 15).

**Movie S7.** Brightfield image stack of 2 h starved dicty cells on a Cy3-labeled, scrambled DNA-conjugated hydrogel under nutrient-free conditions for 5.8 h (2,103 frames, frame interval 10 s, fps 20).

**Movie S8.** Median filtered fluorescence image stack of 2 h starved dicty cells on a Cy3-labeled, scrambled DNA-conjugated hydrogel under nutrient-free conditions for 5.8 h (2,103 frames, frame interval 10 s, fps 20).

**Table S1. DNA sequences used in this paper.** Black text represents the aptamer sequence, blue text represents the poly-T linker.

| **Name** | **Sequence(5’ 🡪 3’)** | **Total Length** |
| --- | --- | --- |
| Anchor DNA sequence | Acrydite/TTTTTGCTTCGGCTCGTATA/BHQ2 | 20 |
| T5 | Cy3/CACCTGGGGGAGTATTGCGGAGGAAGGTTTTTCCAGGTGTATACGAGCCGAAGC | 54 |
| T9 | Cy3/CACCTGGGGGAGTATTGCGGAGGAAGGTTTTTTTTTCCAGGTGTATACGAGCCGAAGC | 58 |
| T33 | Cy3/CACCTGGGGGAGTATTGCGGAGGAAGGTTTTTTTTTTTTTTTTTTTTTTTTTTTTTTTTTCCAGGTGTATACGAGCCGAAGC | 82 |
| Scrambled aptamer | Cy3/GCCTGACGAAGTGAAGGGGCAGGTTGGTTTTTTTTTTTTTTTTTTTTTAAGTCAGGCTATACGAGCCGAAGC | 72 |

**Supporting Figures**

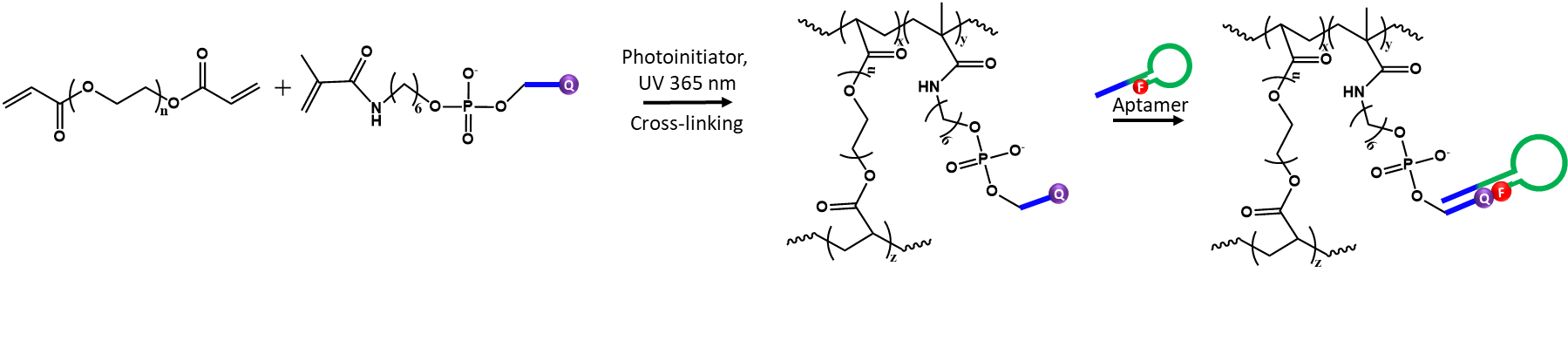

**Figure S1.** Synthesis of the SEMAPHORE hydrogel. Poly(ethylene glycol) diacrylate (PEGDA) and methacrylamide-modified anchor-complementary strands are first coupled via UV photopolymerization, and then the labeled aptamer strands are functionalized onto the hydrogel via hybridization.

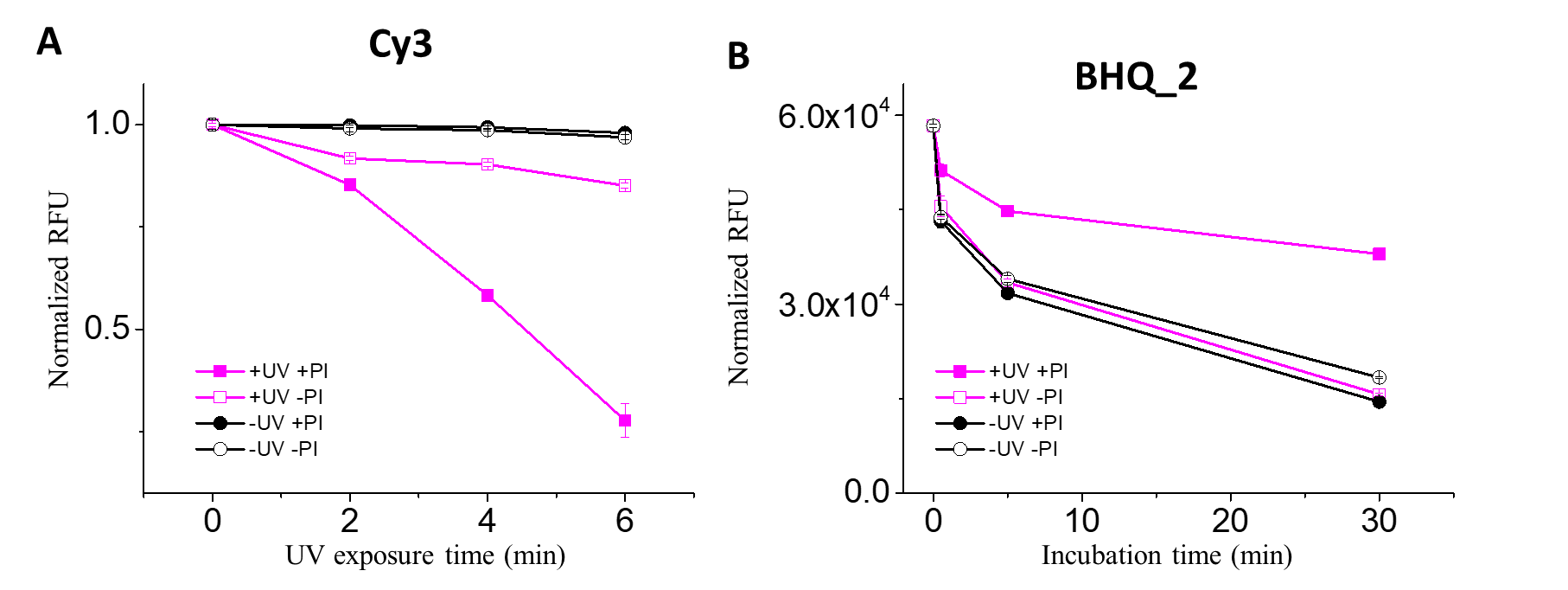

**Figure S2.** Degradation of the fluorophore and dye pair within the aptamer switch design when subjected to different polymerization conditions—with or without ultraviolet (UV), and with or without the free-radical generating photoinitiator (PI)—for hydrogel construction. (A) Measurements of Cy3-labeled DNA alone show that the application of UV, especially in the presence of PI, rapidly degrades the fluorescence. (B) Based on the fluorescence signal from Cy3-labeled strands hybridized to BHQ-2-labeled strands after exposure to combinations of UV and PI. In the presence of UV and PI, the quenching ability of BHQ-2 is substantially reduced.

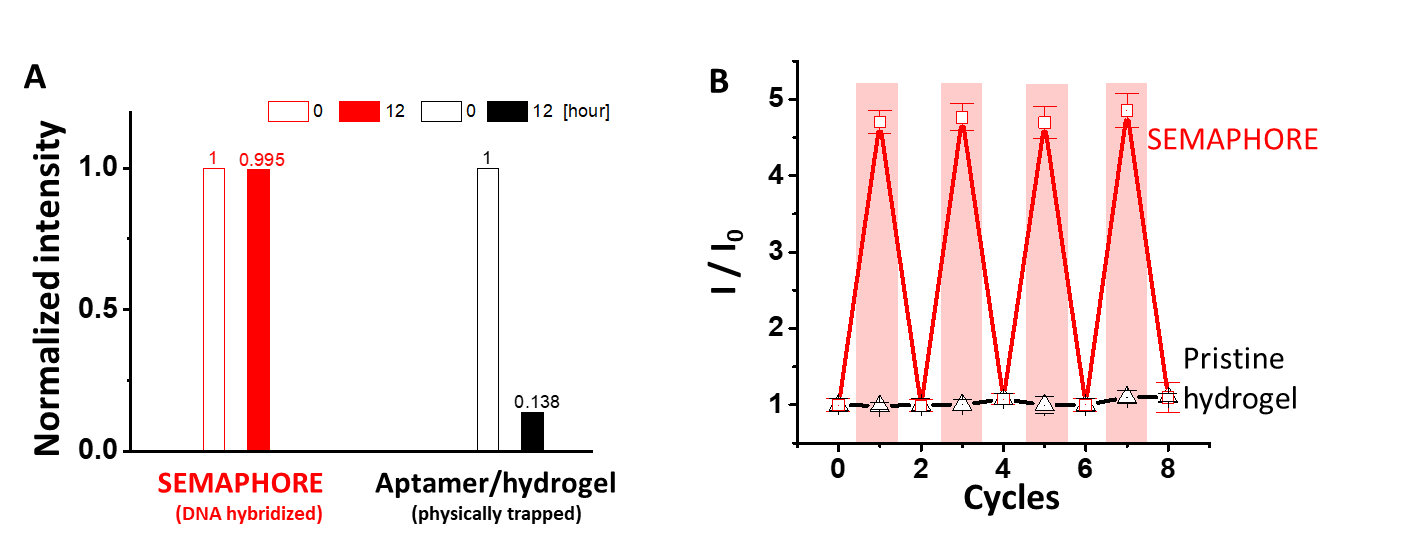

**Figure S3. Long-term stability of SEMAPHORE.** (A) Cy3 fluorescence intensity measured using a fluorescence plate reader before (0 h) and after (12 h) washing with PBS buffer for SEMAPHORE (hydrogel with hybridized aptamers) as compared to a hydrogel where aptamers are trapped only through physical confinement but no hybridization (aptamer/hydrogel). The physically trapped aptamer switches diffused out of the gel during washing, whereas SEMAPHORE showed no fluorescence decrease, indicating stable integration of the aptamer switches. (B) Reversibility test for hydrogel alone (black) compared to the cAMP-responsive SEMAPHORE hydrogel (red). Red shaded regions represent intervals where the gel was exposed to 10 mM cAMP. Each cycle was measured after reaching equilibrium for approximately 1 h. I/I_0_ values were averaged for three independent 1 x 1 mm fluorescence images.

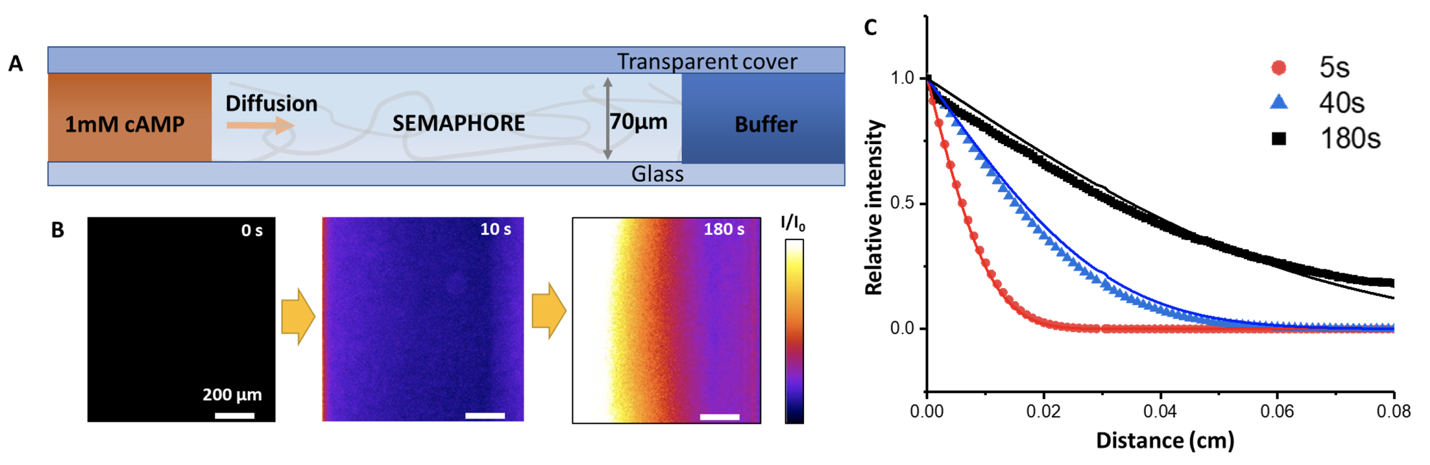

**Figure S4.** Measuring cAMP gradient propagation with SEMAPHORE. (A) Solutions of 0 or 1 mM cAMP were respectively applied to the right and left reservoirs, and fluorescence imaging of SEMAPHORE was used to quantify gradient propagation. (B) Fluorescence images of the SEMAPHORE response to propagation of the cAMP gradient over time. (C) Spatial fluorescence profiles from SEMAPHORE over time show an evolving gradient. Relative fluorescence intensity (I/I_b_) values were averaged from three 1 x 1 mm fluorescence images collected at the center of the hydrogel. Fitting (solid lines) is consistent with a cAMP diffusivity of D = 7.5 cm^2^/s, corresponding well with this molecule’s measured diffusivity in water.

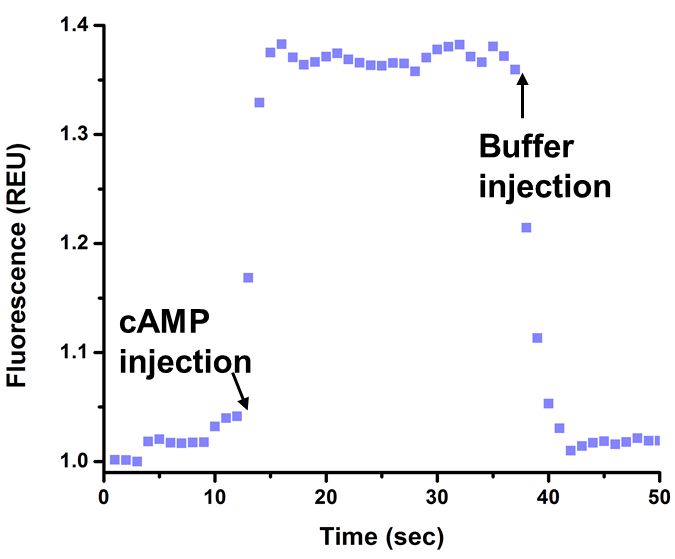

**Figure S5.** Aptamer switch kinetic response to cAMP addition when the T33 aptamer is directly immobilized on a glass surface in the absence of hydrogel. 100 µM cAMP was injected at 13 s and DB was injected at 35 s.

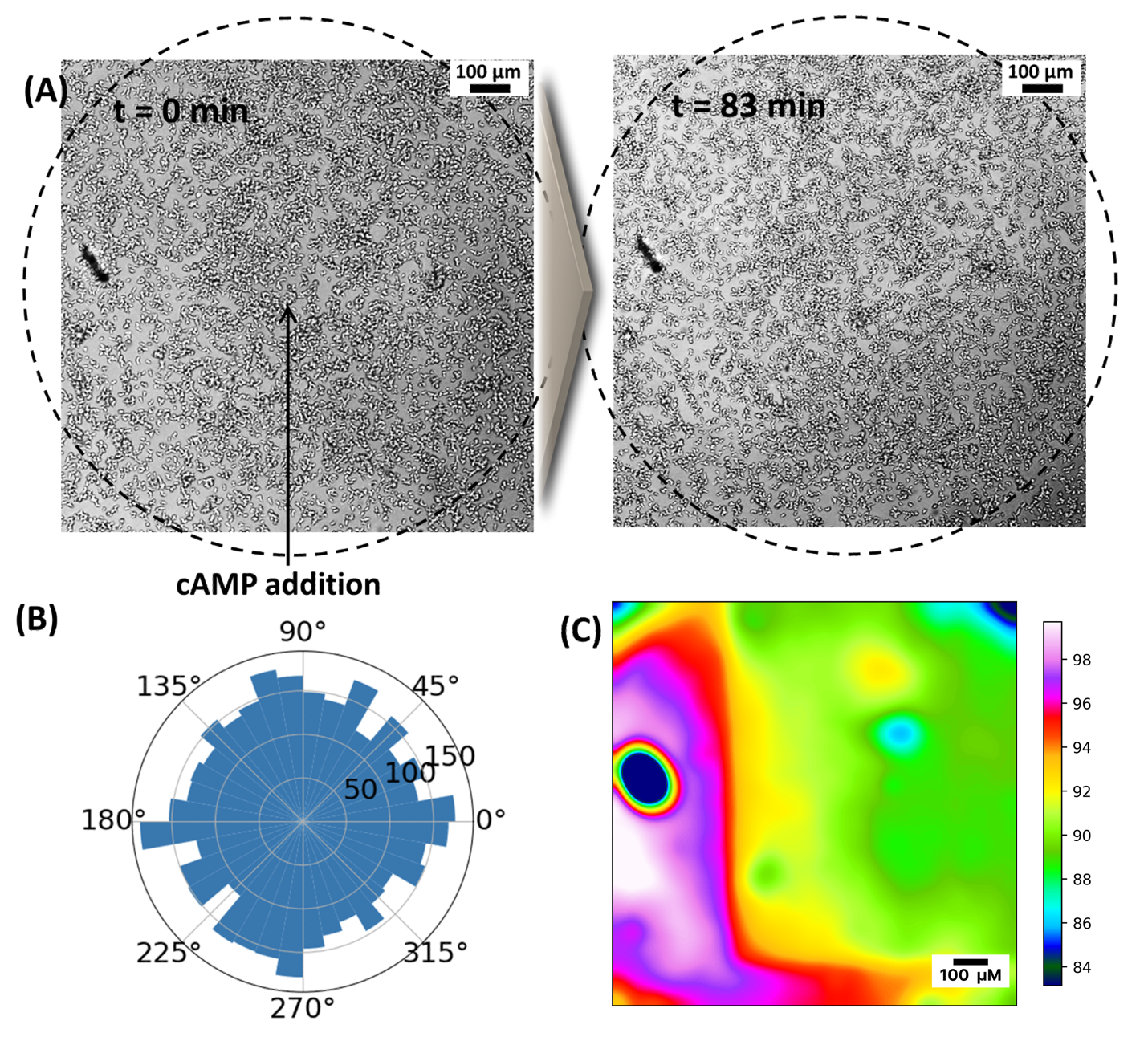

**Figure S6. Monitoring dicty chemotaxis in the absence of a defined cAMP gradient.** (A) Brightfield images at 0 (left) and 83 min (right) after application of 10 µM cAMP across the entire field of view (dotted circle; see **Movie S3**) (B) Polar coordinate histogram of start-to-end cell angular movement for the cells shown above (weighted by velocity). (C) Average fluorescence over the course of 83 min after applying cAMP (see **Movie S4**).

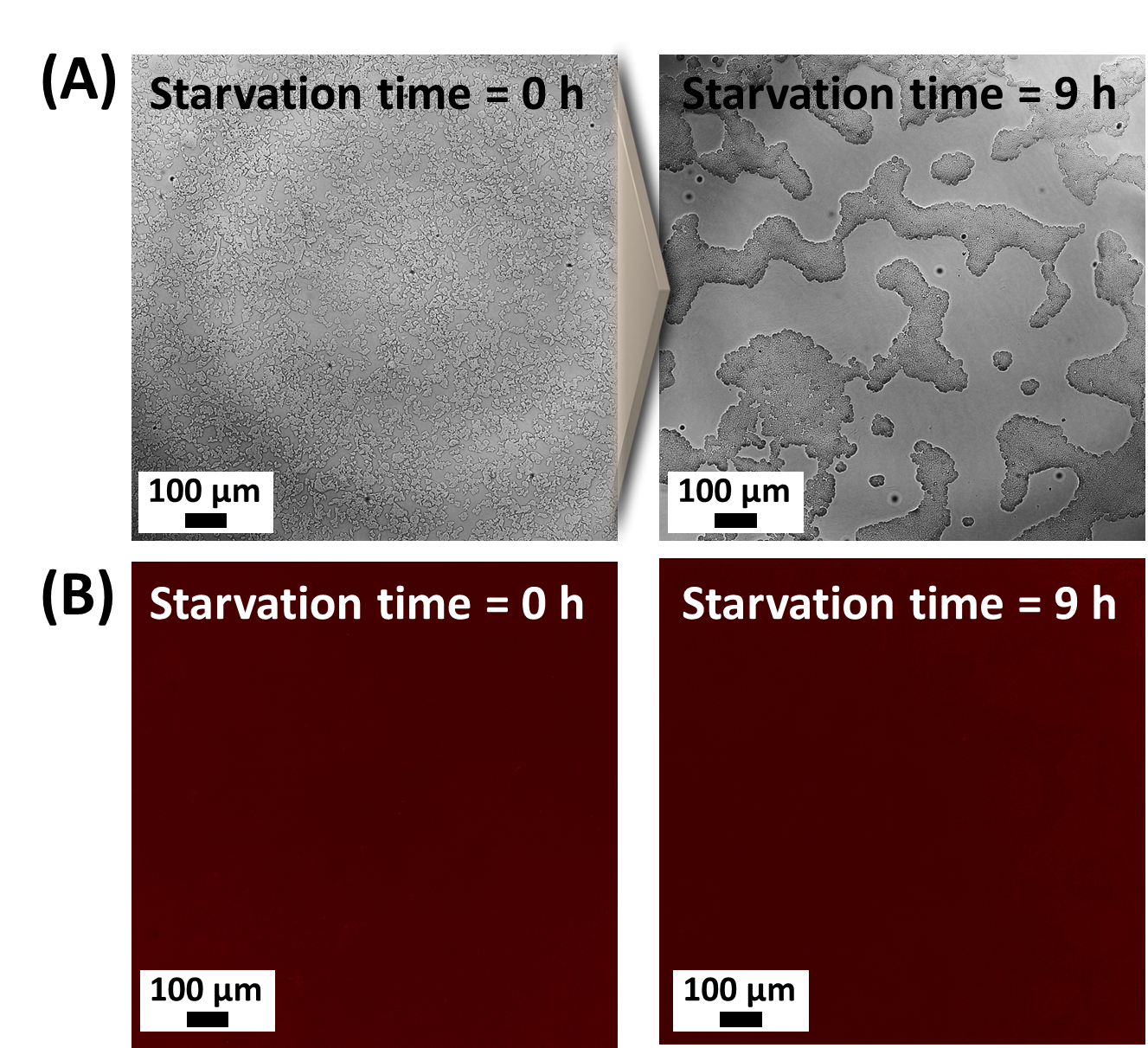

**Figure S7.** (A) Brightfield and (B) fluorescence images of dicty cells in a SEMAPHORE experiment with a scrambled aptamer sequence, which exhibits no response to cAMP secretion.

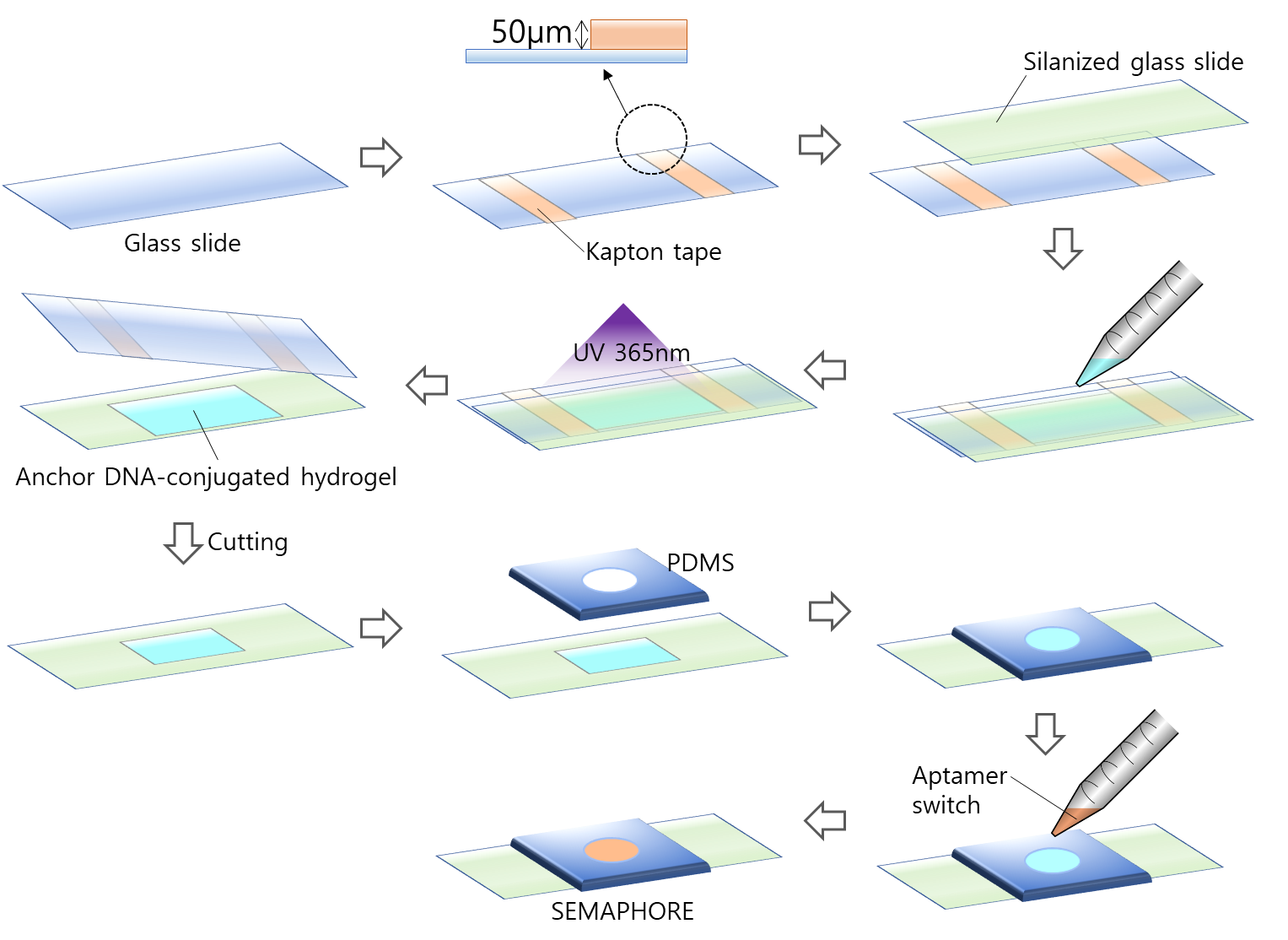

**Figure S8.** Schematic of the fabrication process for the SEMAPHORE hydrogel system within a small PDMS chamber.
